# *Plasmodium falciparum* Acetyl-CoA Synthetase is essential for parasite intraerythrocytic development and chromatin modification

**DOI:** 10.1101/2021.06.13.448207

**Authors:** Isadora Oliveira Prata, Eliana Fernanda Galindo Cubillos, Deibs Barbosa, Joaquim Martins, João Carlos Setubal, Gerhard Wunderlich

## Abstract

The malaria parasite *Plasmodium falciparum* possesses a unique Acetyl-CoA Synthetase (PfACS) which provides acetyl moieties for different metabolic and regulatory cellular pathways. We characterized PfACS and studied its role focusing on epigenetic modifications using the *var* gene family as reporter genes. For this, mutant lines to modulate plasmodial ACS expression by degron-mediated protein degradation or ribozyme induced transcript decay were created. Additionally, an ACS inhibitor was tested for its effectiveness and specificity in interfering with *Pf*ACS. The knockdown of *Pf*ACS or its inhibition led to impaired parasite growth. Decreased levels of *Pf*ACS also led to differential histone acetylation patterns, altered variant gene expression and concomitantly decreased cytoadherence of infected red blood cells containing knocked-down parasites. Further, ChIP analysis revealed the presence of PfACS in many loci in ring stage parasites, underscoring its involvement in the regulation of chromatin. Due to its significant differences to human ACS, *Pf*ACS seems an interesting target for drug development.

## Introduction

Human malaria is caused by protozoan parasites of the genus *Plasmodium* and still affects a huge part of the population living in developing countries located in tropical and subtropical regions. The World Health Organization (WHO) have registered 409,000 malaria deaths in 2019, where *P. falciparum* is responsible for the most severe forms of malaria in humans, causing the majority of deaths (1). Artemisinin Combined Therapy (ACT) combined with the use of long-lasting insecticide-treated bed nets (1) and indoor residual spraying campaigns (2) were successful for the global decrease in malaria infections. Nevertheless, resistance of *P. falciparum* against Artemisinin was reported in Cambodia in 2008 (3), followed by treatment failure along the Thailand-Myanmar border (4) and recently in Central Africa. Therefore, the discovery and development of new effective antimalarial drugs is important to further decrease malaria incidence.

The clinical symptoms of malaria appear during the intraerythrocytic asexual cycle of the parasite. During this process, the parasite exports proteins into the red blood cells (RBCs) (5). Members of the *Plasmodium falciparum* erythrocyte membrane protein 1 (PfEMP1) family are expressed on the RBC surface and they are important virulence factors and relate directly to the persistent nature of the malaria disease (6). PfEMP1-mediated infected erythrocyte sequestration through cytoadherence has several consequences such as the blockage of microvasculature and endothelial activation, the main cause of severe malaria (7). PfEMP1 proteins are encoded by the 45-90-member variant gene family named *var*, and transcriptional switching results in the appearance of different PfEMP1 alleles on the IRBC membrane (8). Expression of PfEMP1 is strictly controlled, and normally only one *var* gene is transcribed (9). Many factors are involved in this process, including histone-modifying enzymes such as acetyltransferases (10) and deacetylases (11, 12), methyltransferases (13), non-coding RNAs (14), chromatin spatial positioning (15) and the presence or absence of HP1 (*Heterochromatin Binding Protein* 1) (16, 17).

Epigenetic marks and their modifiers have been largely studied and their importance in *var* gene expression control has been partly elucidated. The acetylation mark at lysine 9 Histone 3 (H3K9Ac) (18) and the presence of the special histone H2.AZ at promoter regions (19) play a crucial role in *var* gene transcription activation, while the trimethylation of lysine 9 Histone 3 (H3K9me3) and of lysine 36 Histone 3 (H3K36me3) are *var* gene silencing marks (13, 18). *Var* gene expression is semi-conserved so that the same *var* gene is commonly expressed after the next reinvasion (epigenetic memory). A few marks seem crucial for the epigenetic memory, such as di- and trimethylation of lysine 4 Histone 3 (H3K4me2 and H3K4me3), that are related to an active *var* gene locus (20).

Many central metabolites and cofactors are important for the function of chromatin-modifying enzymes and consequently to maintain or alter gene expression regulation in response to alterations in the homeostasis (21). Among them, the cofactor acetyl-CoA was shown to be crucial for protein acetylation in many different organisms and is a substrate for histone acetylation by acetyltransferases (21). In mammalian cells, acetyl-CoA is predominantly synthesized in the mitochondria and the cytosol, generated by the pyruvate dehydrogenase complex (PDH), β-oxidation, or by amino acid transamination that are metabolized in the mitochondrial branched-chain α-ketoacid dehydrogenase (BCKDH) complex (22). Alternatively, in stress conditions, ATP citrate lyase (ACLY) converts citrate in oxaloacetate and acetyl-CoA. Also, Acetyl-CoA Synthetase 2 (ACSS2), uses acetate in an ATP dependent manner to generate acetyl-CoA (23). *P. falciparum* acetyl-CoA generation is different from other organisms. Likewise, acetyl-CoA derived from pyruvate enters the TCA cycle, but, unlike most organisms, most of this cofactor is not generated by PDH. Actually, the absence of PDH, which is apicoplast-localized, generates no parasite growth defect, no alterations in the glycolytic flux and the intracellular acetyl-CoA level (24). On the other hand, *P. falciparum* BCKDH is not capable of amino acid degradation, and, supposedly, through a PDH-like activity, this pathway generates acetyl-CoA from pyruvate (24) and could possibly be a major provider for cellular acetyl-CoA in the parasite.

We characterized Acetyl CoA synthetase of *P. falciparum* as an important source of acetyl-CoA and its knockdown or inhibition led to a loss of epigenetic acetylation marks, resulting not only in parasite growth impairment but also in partial *var* gene silencing. Also, this protein interacts with chromatin, apparently participating in the control of many genes related to parasite metabolism and pathogenesis.

## Results

### PfACS can be tagged and its knockdown leads to decreased PfACS presence

The gene encoding PfACS is localized on chromosome 6 (PF3D7_0627800) and consist of a 4176 bp long gene with an extension of its N-terminus not encountered in human or yeast ACS. In order to monitor PfACS function, a *P. falciparum* NF54 based lineage was created using the plasmid pACS-GFP-HA-DD24-2A-BSDglmS (Supplementary Figure 1). After establishment of the WR99210-resistant lineage, a second round of drug selection with Blasticidin was used to select only PfACS locus-modified parasites (Figure 1A). After initial transfection, parasites started to re-appear in blood smears 15 days after cultivation in the presence of WR99210, selecting for hDHFR cassette containing parasites. Parasites subsequently submitted to Blasticidin treatment became visible in blood smears after 17 days. Correct insertion in the ACS encoding locus was confirmed by site-specific PCRs (Figure 1B).

**Figure 1.**
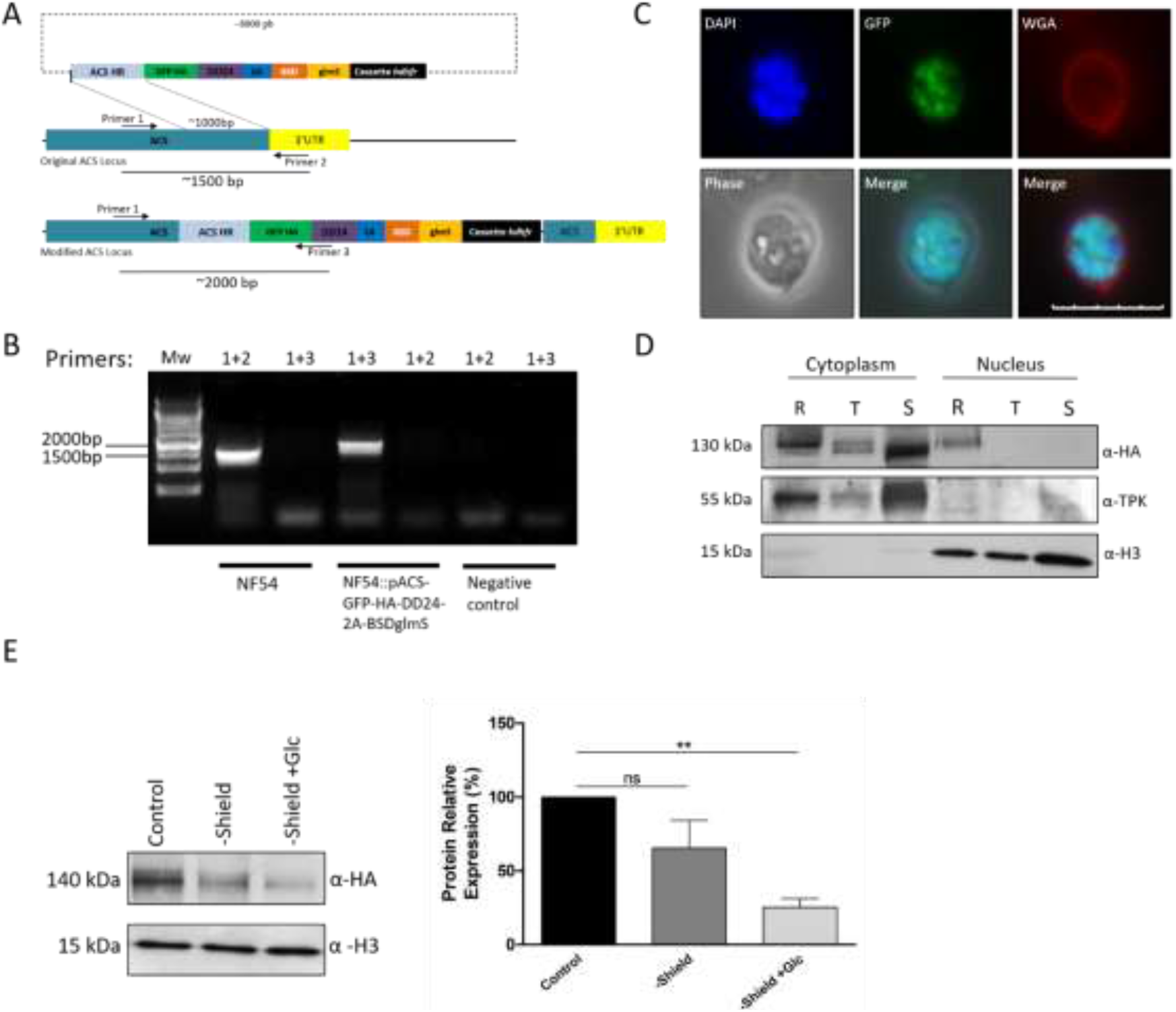
NF54::pACS-GFP-HA-DD24-2A-BSDglmS mutant line construction and conditional knockdown. **A -** Scheme of the plasmid used in the generation of the NF54::pACS-GFP-HA-DD24-2A-BSDglmS mutant line indicating the integration locus for Acetyl-CoA Synthetase (PF3D7_0627800). **B -** PCR reaction using primers designed for amplification of the mutated locus (primers 1+3) and the WT NF54 locus (primers 1+2), which was absent in the mutant line. **C –** Fluorescence microscopy of a schizont of the transgenic line exhibiting GFP expression. DAPI was used as parasite nuclear marker and WGA as erythrocyte surface marker. Scale bar: 10 µm. **D –** Western blotting for PfACS subcellular localization, indicating its presence in cytoplasmic fractions in all stages and in the nuclear fraction in ring stage. **E –** Western blotting after 24h PfACS knockdown with Shield withdrawal (35% protein reduction) and Shield withdrawal plus 2.5 mM Glucosamine (75% of protein reduction). Bars indicate mean ± SD. Statistical test: Student t-test **P<0.01, ns= non-significant. Values from three independent experiments are shown.

The correct integration generated an ACS protein fused to GFP and HA-tag, which allowed GFP tagged ACS visualization by fluorescence microscopy. GFP expression was easily observed in ring (Supplementary Figure 2), trophozoite (Supplementary Figure 3) and schizont stages (Figure 1C), indicating the presence of this protein throughout the intraerythrocytic asexual cycle of the parasite. To localize PfACS, NF54::pACS-GFP-HA-DD24-2A-BSDglmS parasites were separated in cytosolic and nuclear fractions and submitted to western blot analysis using Histone H3-specific antibodies to mark nuclear fractions and Thiamine Pyrophosphokinase (TPK)-specific antibodies to mark the cytoplasmic compartment (Figure 1D). While PfACS was detected in the nuclear and the cytosolic fraction in ring stage parasites, it was no longer present in nuclei in trophozoite and schizont stages.

In order to evaluate the effect of PfACS knockdown, the mutant line was split into three groups: a control group, continuously grown in the presence of Shield, -Shield group, where Shield was omitted from the culture medium, and the third knockdown group, where besides Shield removal, 2.5 mM Glucosamine was added to the culture medium (-Shield+Glc group). The PfACS protein decreased in both knockdown groups compared to the control, but the combination of transcript ribozyme-mediated knockdown and Shield removal was more efficient compared with the second group (Figure 1E). We observed a significant 75% PfACS decrease by inducing knockdown of transcript and protein.

### PfACS knockdown or inhibition impairs parasite growth and results in decreased histone acetylation

During PfACS knockdown, parasite viability was monitored during 72h of treatment, starting from ring stage synchronous parasites (0h) (Figure 2A). Parasites from both knockdown groups appeared unhealthy, starting after 24h of treatment, when synchronicity was partially lost compared with the control group. After 48h, no healthy parasites could be found and, in the following days, blood smears from the -Shield+Glc group contained almost no parasites. Although remaining parasites from the -Shield group could still be seen in the following days, apparently capable of replication and reinvasion, they appeared amorphous. This indicates that PfACS is an important protein for parasite survival, since a 35% decrease was sufficient to interfere with the proper development of the parasite. A similar assay was performed in *P. falciparum* NF54 WT. Instead of knockdown, a specific ACS inhibitor (iACS) successfully tested in human (25) and mouse cells (26) was used. We cultivated parasites in the presence of iACS concentrations ranging from 30 μM to 50 μM and documented the phenotype observed throughout 72h (Figure 2A). Similar to the knockdown experiment, parasites lost synchronicity after 24h, appearing still viable in 30 μM and very unhealthy in higher concentrations of iACS. After that, parasites became unviable in an inhibitor dosage-dependent manner.

**Figure 2.**
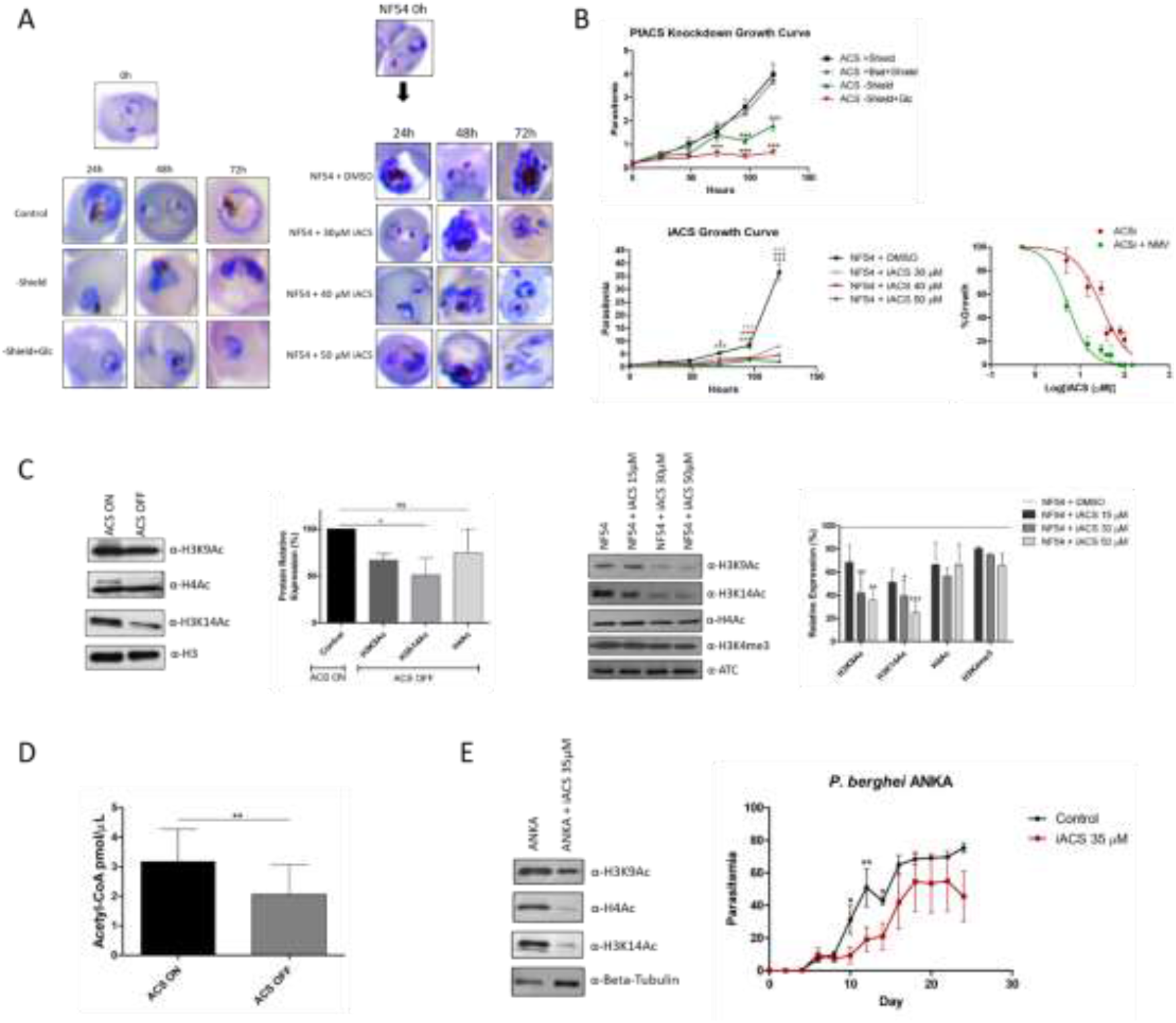
ACS inhibition or knockdown leads to growth defect and loss of epigenetic marks. **A –** Left panel indicates the mutant NF54::pACS-GFP-HA-DD24-2A-BSDglmS phenotype after ACS conditional knockdown through Shield withdrawal and Shield withdrawal in the presence of Glucosamine, where parasites were found asynchronous after 24h, very unhealthy after 48h and amorphous and unviable after 72h in both conditions compared to the controls. The right panel shows WT NF54 parasites treated with iACS, which induced the same phenotype observed in PfACS knockdown parasites when the dpsage was increased. **B –** At the top, the *in vitro* growth curve indicates that PfACS knockdown through Shield withdrawal and Shield withdrawal in the presence of 2.5mM Glucosamine impairs *P. falciparum* growth. At the bottom on the left, the presence of iACS also generates a growth defect in *P. falciparum* cultures in a dose dependent manner (For this experiment, cultures were diluted and virtual parasitemias are shown). At the right, the *in vitro* IC50 curve of iACS is shown. iACS IC50 decreased from 30.14 to 5.49 μM using NMVs loaded with iACS compared to iACS alone. **C –** Western blotting exhibiting the loss of epigenetic marks in NF54::pACS-GFP-HA-DD24-2A-BSDglmS parasites after 24h PfACS knockdown and NF54 treated 24h with iACS, where the marks H3K9Ac and H3K14Ac in both conditions were significantly decreased. **D -** Acetyl-CoA cellular levels were significantly affected by PfACS knockdown. **E -** Western blotting shows a strong depletion of H3K9Ac, H4Ac and H3K14Ac acetylation epigenetic marks in *P. berghei* after 18h of iACS treatment. On the right, curve of mice infected with *P. berghei* and treated with 35 μM iACS during the first 3 days of infection compared to control mice, treated with DMSO. A delay in parasite growth compared to the control group can be observed, where parasitemia became undetectable in one of 5 mice. Bars indicate mean ± SD. Statistical tests: (B) and (E) Two-way ANOVA with Bonferroni’s post-test. (C) and (D) Student t-test. *P<0.05, **P<0.01, ***P<0.001.

Parasite viability was then evaluated by a 120h growth curve during PfACS knockdown (Figure 2B). For that, the mutant parasite line was synchronized in ring stage (0h) and the knockdown groups were separated as described above. After 72h, the ability to grow was significantly affected in -Shield+Glc parasites compared with the control groups. After 96h of treatment both knockdown groups were unable to grow. As expected, due to the lower knockdown efficiency in the -Shield group, these parasites were not completely affected at the first days of the growth curve, reaching a maximum parasitemia of 2% at the end of the experiment, approximately half of the control group’s parasitemia. The -Shield+Glc group growth was completely impaired by PfACS knockdown, pointing to a significant function of this protein in parasite growth and development.

A similar growth assay was performed varying iACS concentrations from 30 to 50 μM and using NF54 WT parasites treated with DMSO as the control group (Figure 2B). Parasitemia from iACS-treated groups started to slightly decrease after 48h and, after 72h, parasites treated with 40 and 50 μM iACS expanded significantly less than the control group. After 96h, all iACS-treated groups grew significantly less than the control group, indicating that iACS led to parasite growth impairment in a dose-dependent manner. We determined the *in vitro* iACS IC_50_ and, in order to improve inhibitor effectiveness, we encapsulated the compound in nano-multilamellar vesicles (NMV) stabilized by interlayer hydrogen bonds (27) (Figure 2B). iACS IC_50_ decreased from 30.14 to 5.49 μM using NMVs loaded with iACS compared to iACS alone, a 5.5-fold reduction that can be observed in the IC_50_ curves.

Considering that a knockdown induced by Shield-1 withdrawal and Glc addition was most effective, the next experiments were carried out using this method where the control group is referred to as ACS ON and the knockdown group as ACS OFF. In order to show an effect of PfACS depletion/inhibition also on histone modification, we tested the effects of PfACS knockdown on the parasite’s H3K9Ac, H3K14Ac and H4Ac histone acetylation marks (Figure 2C). Western blots of total extracts indicated that H3K9Ac was reduced by 40% compared with the control, while H3K14Ac decreased 50%. In contrast, the H4Ac mark was not significantly affected by ACS knockdown. We also tested the effects of iACS on the same parasite acetylation marks plus one trimethylation mark, H3K4me3. The total amount of H3K9Ac and H3K14Ac marks was significantly reduced in a dose dependent manner in dependent on the presence of iACS, compared with the vehicle-treated control (Figure 2C). As expected, H3K4me3 was not affected by the treatment. The iACS presence induced a stronger decrease of the same acetylation marks compared with the PfACS knocked-down mutant line. This suggests that iACS has a specific activity against PfACS protein, however, other off-target effects are also possible.

In order to measure whether acetyl-CoA parasite cellular levels would be affected by ACS knockdown, we measured acetyl-CoA levels using a fluorescent quencher. Upon knockdown, there was significantly less acetyl-CoA present (Figure 2D), further pointing to the view that this protein is an important for the production of acetyl-CoA in *P. falciparum*.

ACS is conserved in other *Plasmodium* species. Given the high similarity between *Plasmodium berghei* Acetyl-CoA Synthetase (PbACS; PBANKA_1126500) and PfACS, we tested whether iACS would be similarly effective against *P. berghei* in the murine model. For this, we initially tested if histone acetylation marks would be affected by treatment with iACS at a final concentration of 35 μM. A clear decrease in all acetylation marks tested was observed (Figure 2E). Interestingly, in contrast to immunoblots using material from *P. falciparum*, the acetylation mark H4Ac also seemed strongly reduced in *P. berghei* after iACS treatment. However, this experiment was not done in triplicate. Subsequently, we infected mice with *P. berghei* WT parasites and treated the control group with DMSO and the treatment group with 35 μM iACS for the first three days after infections and followed parasitemia every two days for 24 days (Figure 2E). After day 8 post infection, iACS treated parasites were not growing as fast as the control group. Interestingly, after day 6, one mouse of five from the iACS-treated group had undetectable parasite burden until the end of experiment. iACS was effective in impairing *P. berghei* parasite growth as observed in *P. falciparum*, considering that parasitemias from treatment and the following days were significantly affected in relation to the control.

### PfACS knockdown or inhibition leads to decreased *var* gene transcript levels

To further support the hypothesis that the decreased presence of PfACS not only leads to metabolic interference of parasite growth but also interferes in epigenetic modifications resulting in an alteration of transcription/transcript levels, we used *P. falciparum var* genes as a surrogate marker. Acetylated H3K9 was described as the mark for actively transcribed *var* loci, therefore, short time inactivation or decrease of PfACS should lead to decreased *var* transcript levels. To test this, PfACS was knocked down or inhibited for a period of ∼20h, which is before parasitemia starts to decline due to treatment, and performed *var* gene transcript analysis. This was done on previously CHO-CD36-cytoadherence-selected mutant parasite lines, which are supposed to transcribe preferentially determined *var* genes. Western blotting was used to confirm that protein became expressed after treatment end (Supplementary Figure 4). After two panning assays, the most expressed *var* genes were PF3D7_1240600 and PF3D7_0412400, which encode CD36 binding PfEMP1s (28, 29), and PF3D7_0420900 (Figure 3). Under PfACS knockdown (Figure 3A) and iACS treatment (Figure 3B), a very similar *var* transcript profile for the two different treatments was observed, however, the transcripts were observed in relatively smaller quantities. The transcript of the most expressed *var* gene, PF3D7_1240600, decreased 2.3-fold during knockdown and 3-fold during iACS treatment. PF3D7_0412400 was reduced 4.5-fold for both knockdown and iACS treatments, and PF3D7_0420900 decreased by 2.5-fold during PfACS knockdown, followed by 2-fold change during iACS treatment. The transcript quantity reductions were statistically significant (P<0.001), indicating that PfACS is involved in the epigenetic modification leading to *var* gene locus activation.

**Figure 3.**
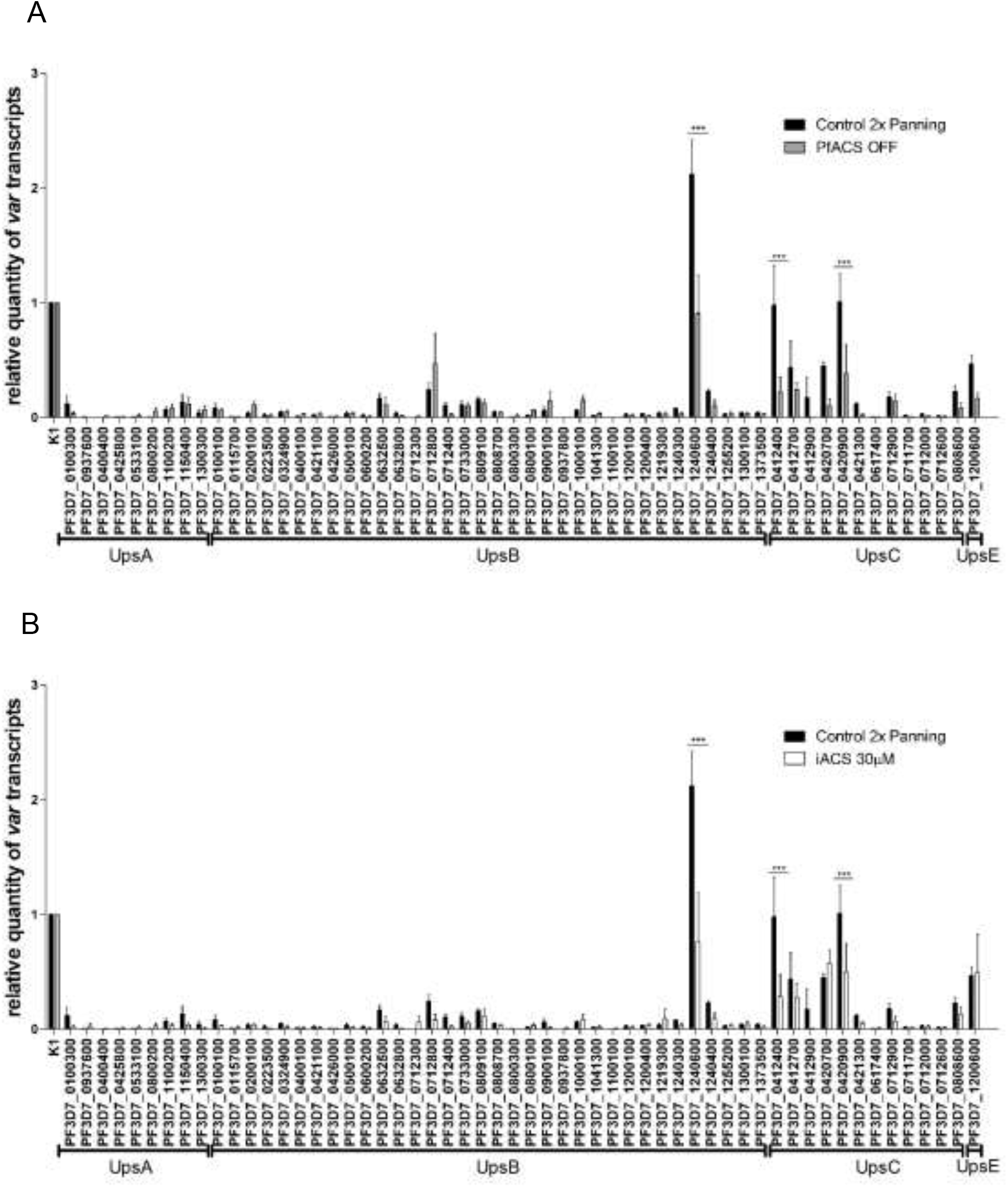
PfACS knockdown or inhibition leads to *var* gene de-activation. *var* gene transcript profiles exhibiting PfACS knockdown (Top) and iACS 30 µM treated (Bottom) gene expression compared to controls. The de-activation of the three control selected genes can be observed in both conditions, being a statistically significant (P<0.0001) reduction. Bars indicate mean ± SEM. Statistical test: Two-Way ANOVA with Bonferroni’s post-test. ***P<0.001. These results are from biological triplicates.

A recent study indicated the physical presence of PfACS at *var* gene loci (30). To test if PfACS has also a role in the epigenetic memory of *var* gene transcription, we induced protein knockdown or inhibition for one reinvasion cycle and then treatments were interrupted to allow normal PfACS protein levels/normal PfACS activity. Parasites in which PfACS had been re-stabilized (RE-ON), an increase in the quantity of PF3D7_1240600, PF3D7_0412400, PF3D7_0632500 and PF3D7_0420900 was observed. Of note, these are the same transcripts that were present before the knockdown/treatment, and they were in significantly higher levels than in the untreated controls (Figure 4A). In iACS treated and recovered parasites, a significant increase in the transcript levels of PF3D7_0420900 compared with the control was observed (Figure 4B). Thus, parasites maintained the expression of basically the same *var* genes after one PfACS knockdown or inhibition cycle, indicating that this protein does not play a role in the coordination of the epigenetic memory of *var* transcription.

**Figure 4.**
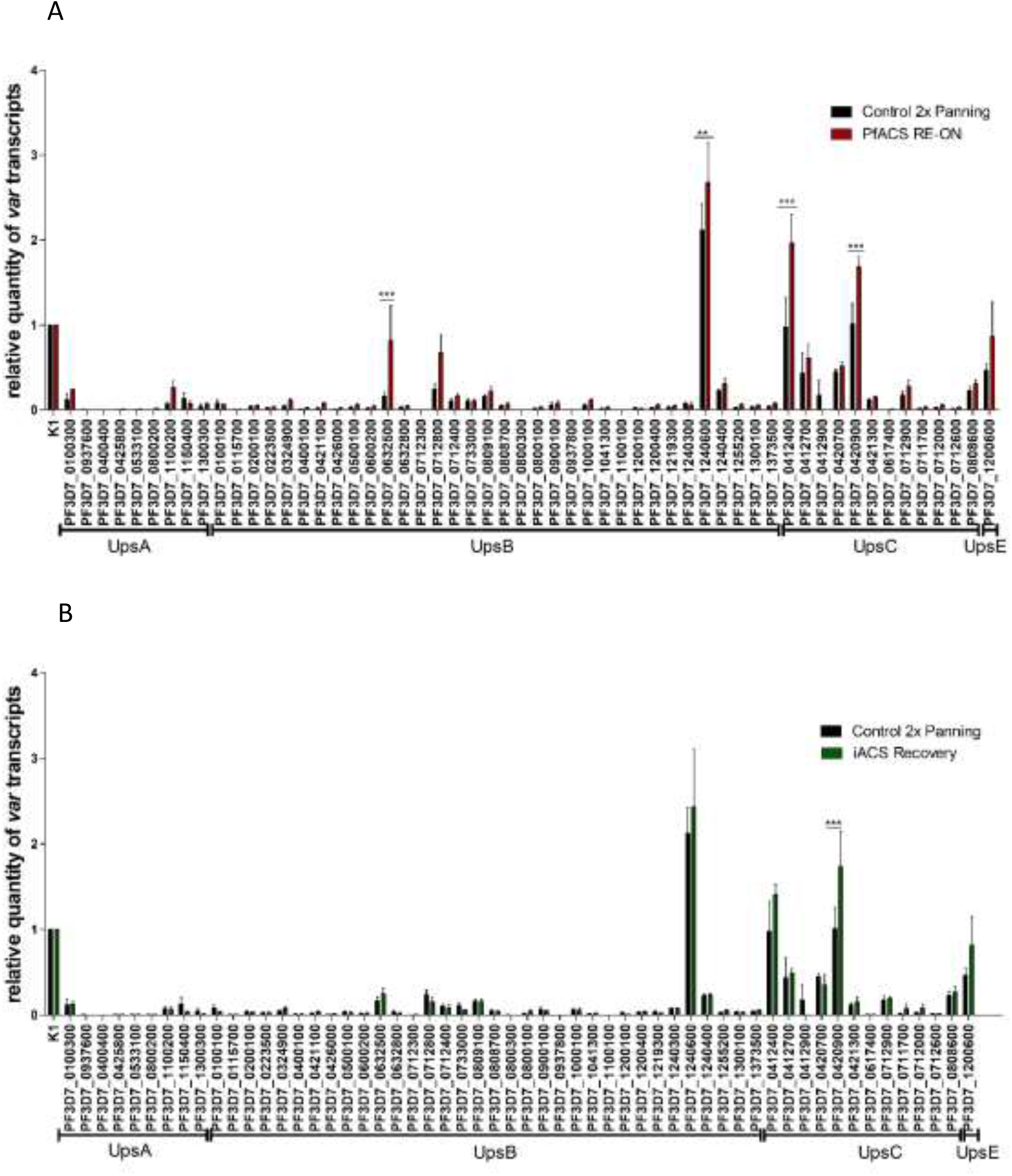
PfACS knockdown or inhibition does not affect the transcriptional memory of *var* genes. **A -** ACS RE-ON parasites, where PfACS knockdown was induced for one reinvasion cycle and re-established after a second reinvasion cycle, had their *var* transcript profile evaluated. The same set of genes as observed in the control, except for PF3D7_0632500, showed relatively more transcripts than the never treated control. PF3D7_0632500 was found inactive before and during PfACS knockdown, starting to be expressed significantly higher compared to control after protein recovery. **B -** iACS treated and recovered parasites, where PfACS inhibition was induced for one reinvasion cycle after which iACS was removed from the medium, permitting normal PfACS function in the subsequent second reinvasion cycle. *Var* transcript analysis showed that the same set of transcripts as observed in controls had their levels restored. Of note, PF3D7_0420900 showed significantly higher levels than the control. **P<0.01, ***P<0.001. Bars indicate mean ± SEM. Statistical test: Two-Way ANOVA with Bonferroni’s post-test. The experiments were done in biological triplicates.

### PfACS knockdown or inhibition interferes in IRBC cytoadherence

The parasite line containing a tagged PfACS, selected for cytoadherence on CHO-CD36, actively transcribed *var* genes encoding for CD36-binding PfEMP1s. To elucidate whether a temporary PfACS knockdown influenced IRBC cytoadherence, we performed cytoadherence assays with confluent CHO cells expressing the human CD36 receptor (31), the same cell lineage used for panning assays. Then, PfACS knockdown or inhibition was induced. In the following, the cytoadhesion ratio (number of iRBCs per CHO-CD36 cells) was compared with untreated control cultures (Figure 5B) by using phase contrast images from adhered parasites (Figure 5A). During knockdown or iACS treatment, parasites did not adhere to CHO-CD36 cells in the same number as the control parasites. Apparently, the decreased presence of PfACS and consequently the decreased abundance of *var* transcripts also led to decreased cytoadherence, probably by diminished PfEMP1 presence. We then performed cytoadherence assays with ACS RE-ON and iACS recovered parasites to test whether the adherence phenotype would be similar to the one before PfACS depletion/inhibition (Figure 5A). Interestingly, we noticed that the cytoadherent phenotype became more prominent in iACS recovered parasites, where the adherence ratio was significantly higher than in the controls (Figure 5B). The cytoadherence pattern of PfACS-knocked down and re-established (RE-ON) parasites was similar to the one of the control cultures. Taken together, PfACS is important for *var* gene activation and subsequently also for cytoadherence of infected red blood cells, although PfACS depletion does not affect the transcriptional memory of *var* genes.

**Figure 5.**
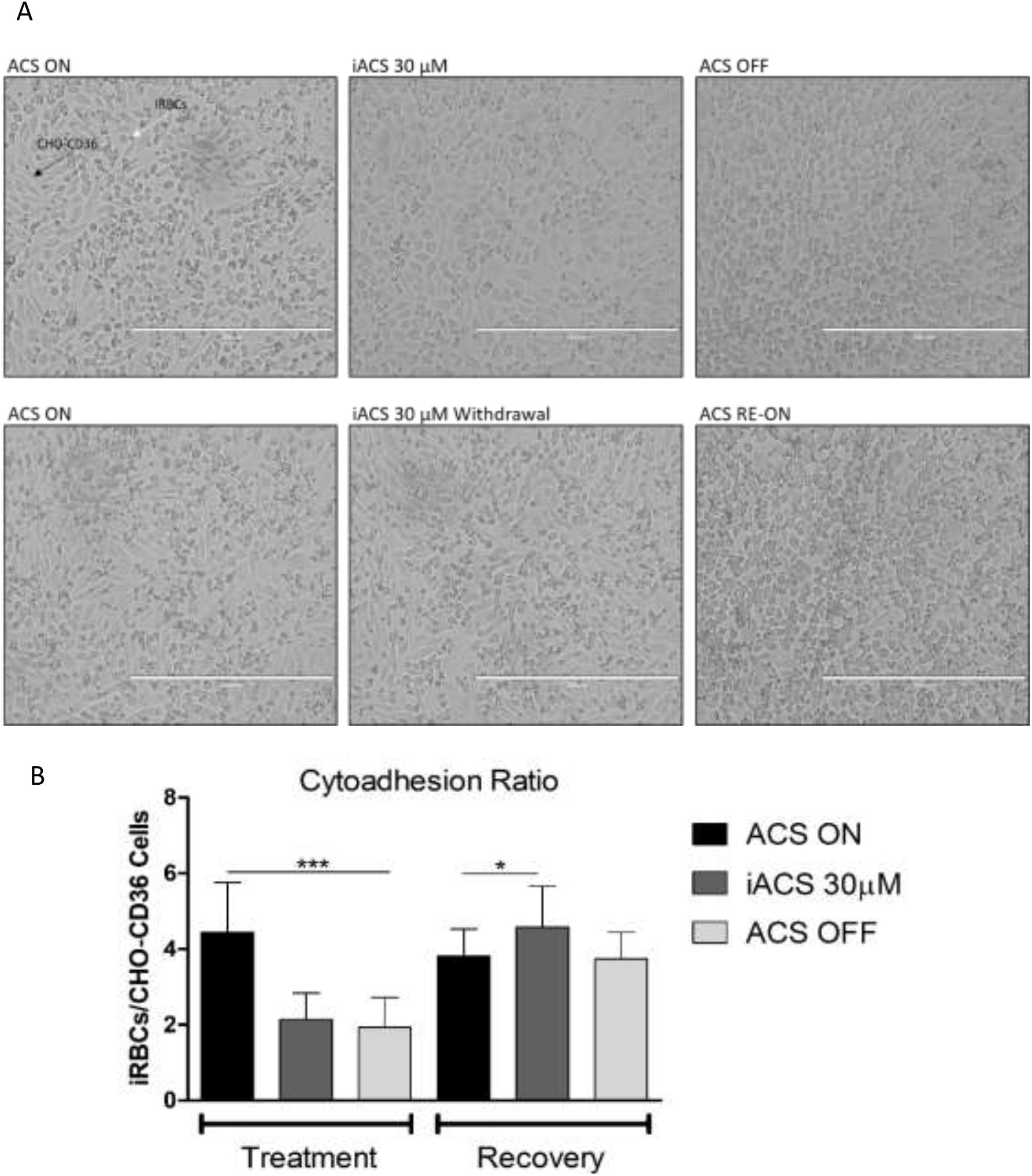
PfACS depletion reverts cytoadherent phenotype. **A -** Phase contrast microscopy of CHO-CD36 binding iRBCs during PfACS depletion (Top) and recovery (Bottom). Images exhibit iRBCs interacting with CHO-CD36 cells (black arrows), where the small darker particles are IRBCs (white Arrows). Control parasites are more cytoadherent (ACS-ON - left), while iACS treated (middle) and ACS OFF (right) lost their ability to adhere compared to controls. After PfACS knockdown release, cytoadherence phenotypes were re-established and parasites could again adhere to CHO-CD36 cells. **B -** Mutant parasites cytoadherent to CHO-CD36 submitted to PfACS knockdown or iACS mediated inhibition had their phenotype significantly decreased compared to the control. After re-establishment of PfACS, parasites regained their cytoadherent phenotype, and iACS treatment recovered parasites became more adherent than before iACS treatment. Bars indicate mean ± SD. Statistical test: Student t-test ***P<0.001, *P<0.05.

### PfACS interacts with chromatin

Given that PfACS interferes with histone acetylation and co-precipitated with *var* intron sequences (30), we tested to which gene loci PfACS was recruited in ChIPseq assays. For that, ring stage parasites were harvested, and PfACS was immunoprecipitated using an HA antibody and sequenced the DNA present in the samples. We found that PfACS interacts with many different regions across the parasite genome (Supplementary Table 1) and peak calling of enriched loci returned a total of 3002 different gene loci that are expressed in all stages of the parasite intraerythrocytic cycle. PfACS was found primarily surrounding exonic regions and much less in intronic and intergenic regions. When intronic regions were identified, these comprised pseudogenes. In general, PfACS exhibited mainly a non-telomeric distribution. The sub-telomeric genes found enriched in our analysis belong mainly to virulence-associated variant genes, such as *rif, stevor, surf* and *var* genes.

For an overview, all genes with an arbitrarily set qvalue < 5.03×10^−16^ across all chromosomes were plotted. PfACS seemed to recruit to loci in all 14 chromosomes, mostly in non-telomeric regions, although telomeric genes were also represented (Figure 6A; Supplementary Table 2). When genes obtained from the peak calling analysis were separated into Panther Protein Class (Gene Ontology (GO)/Panther (http://geneontology.org/)), we observed that 21.9% of the database-mapped proteins belong to nucleic acid metabolism proteins, which include Helicases, Transcription factors and RNA binding proteins (Figure 6B). The second group included metabolite interconversion enzymes, composed of proteins related to the central metabolism such as Acyl-CoA Synthetase, Citrate Synthase, and Pyruvate Dehydrogenase. The third most prominent protein class included protein modifying enzymes (chaperones and different protein kinases). In this analysis, enzymes were identified that are essential for cell cycle maintenance and which are usually regulated by the state of their surrounding chromatin (32). Probably, the presence of PfACS near chromatin works as an important source of Acetyl-CoA for histone acetylation. Interestingly, among the relatively small number of promoter regions significantly enriched in our analysis, we found a peak for one *var* gene encoding PfEMP1 PF3D7_0800100 (P=1.29⨯10^−37^) (Figure 6C). This corroborates results shown above that point to a role of PfACS in *var* gene transcription. Among multigenic families, we observed significant enrichment of PfACS at regions covering 8 members of the variant family *surf* genes in non-telomeric regions, 12 genes that belong to the FIKK kinase family (*fikk*), 38 members of the family *Plasmodium* Helical Interspersed Sub-Telomeric (PHIST) protein in non-telomeric and sub-telomeric regions, 15 members of the STEVOR family, 5 Pfmc-2TM members, 7 *var* genes, 11 *rif* family members, and 11 Acyl-CoA protein family members, represented in a section graph (Figure 6D). This indicates that PfACS binds chromatin all over the parasite genome and therefore is possibly part of a gene expression regulator complex. When analyzing the Gene Ontology groups of the ChIPseq-identified gene loci by their significant enrichment, we found that two groups were enriched in the cellular component groups, 26 in the molecular function group and three in the biological process groups (Supplementary Table 4). The enrichment was concentrated in GO groups encompassing genes encoding proteins with regulatory function in the nucleus.

**Figure 6.**
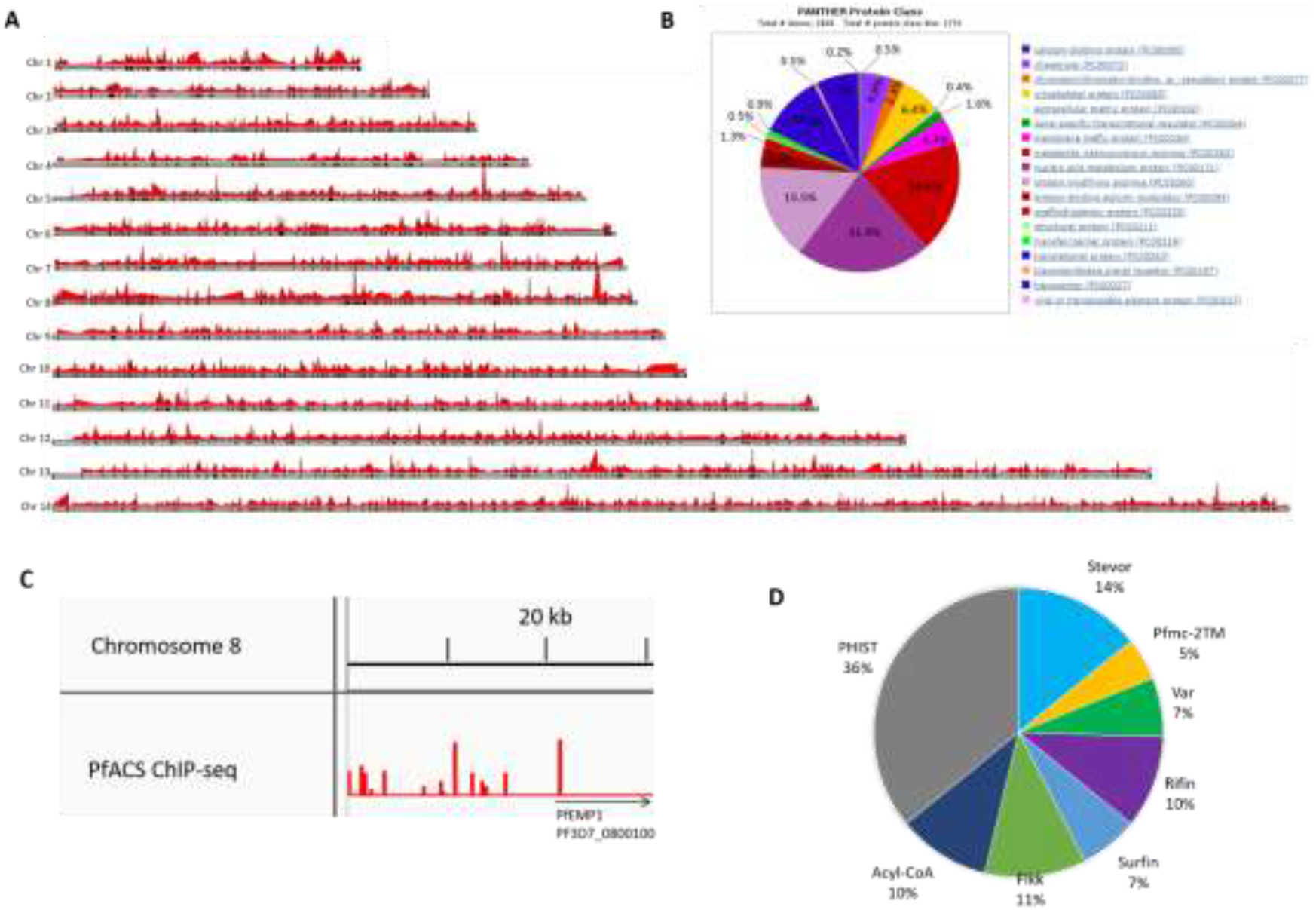
PfACS interacts with chromatin. **A -** All genes with qvalues < 5.03×10^−16^ obtained from PfACS ChIP-seq peak calling across parasite chromosomes are marked by black bars. **B -** GO/Panther Protein Class section plot of PfACS ChIP-seq enriched genes. Peak calling results generated a total of 3002 genes, 2839 found in GO/Panther databank. 1276 hits for GO/Panther Protein class were obtained and analysis indicated that 21.9% of it belong to Nucleic acid metabolism involved proteins. **C -** PfACS ChIP-seq peak of the PfEMP1 PF3D7_0800100 (P=1.29⨯10^−37^). The peak is located at the promoter region of this *var* gene, located at the chromosome 8. **D –** Section plot that represents the different gene families identified in the analysis.

## Discussion

Acetyl-CoA synthetase is a key enzyme in many organisms and acts at a central point of the energy metabolism. Many higher eucaryotes possess different forms of ACS which either contribute for the energy metabolism or provide acetyl moieties for protein modification, including chromatin modification. In the plasmodial genome, only one ACS was identified, indicating that the protein must exert its activity both in the nucleus and the cytoplasm. PfACS seemed to act not only as a metabolic enzyme but also as a player in epigenetic regulation, probably co-acting with histone acetyltransferases. To characterize this enzyme, a mutant line was established that allowed for its knockdown. Additionally, an inhibitor that was previously used to inhibit ACSS2 in mouse cells (26) was also tested. Previous work had revealed that the inhibitor also possesses anti-cancer activity, once it showed to reduce tumors in two different mouse models (25, 33). This was due to its effectiveness inhibiting ACSS2 and consequently reducing cellular acetyl-CoA, which is important for tumor growth. Accordingly, the inhibitor also interfered with *P. falciparum* growth and PfACS function.

Here, a knockdown system that allows transcript depletion of the gene of interest at the same time as its protein destabilization was employed. The architecture resembles that of the knock-sideways approach (34), with the difference that instead of using 2xFKBP domains for mislocating, a degron with a slightly extended knockdown amplitude (DD24 (35)) and an additional glmS ribozyme was used. With both regulons activated, a 75% protein depletion was achieved. Through fusion to GFP, PfACS became detectable in ring, trophozoite and schizont stages, although the subcellular localization was not determinable by microscopy. Subcellular fractioning demonstrated an unusual localization where PfACS is initially nuclear and cytoplasmatic in ring stage, but then exclusively cytoplasmatic in trophozoites and schizonts. Considering that acetyl-CoA diffuses freely through nuclear pores (36, 37), trophozoite and schizont protein acetylation may continue normally in these stages. One hypothesis for the changing presence of PfACS in the nucleus from ring to trophozoite stage is that in ring stage there is a high demand of acetyls for epigenetic programming, necessary for subsequent stages of the parasite. In ring stage, the epigenetic control of different multigenic families, including *var* genes, is important for the subsequent stages of the parasite.

In mammalian cells, ACSS2 is a dynamic protein that migrates from the cytoplasm to the nucleus and vice-versa according to stress conditions such as hypoxia (38). In normal cellular conditions, ACLY is an important source of acetyl-CoA in both cytoplasm and nucleus in most cells. However, in stress conditions such as glucose and oxygen deprivation, ACSS2 migrates from the cytoplasm to the nucleus and becomes the main acetyl-CoA supplier for histone acetylation. These stress conditions are the typical environment of cancer cells, which makes ACSS2 a prime target for cancer drug development (25, 33). Li and colleagues demonstrated that in human glioblastomas, glucose deprivation induces AMP-activated Protein Kinase (AMPK) to phosphorylate ACSS2, which exposes its nuclear localization signal and allows protein migration from the cytoplasm to the nucleus through importin (39). AMPK has not been found in *P. falciparum*, but the putative and uncharacterized SNF1-related serine/threonine protein kinase KIN (PfKIN; PF3D7_1454300), identified by BLAST at PlasmoDB perhaps acts in the AMPK signaling pathways. Interestingly, this protein is present in our PfACS ChIP-seq peak calling results, which indicates that its transcription may somehow be regulated by PfACS. The phenomenon that PfACS apparently is no longer able to enter the nucleus in trophozoites and schizonts is worth further exploration. Of note, the RNA exosome related factor Rrp44 is also present in the nucleus and the cytoplasm in ring stage parasites but exclusively in the cytosol in later forms (40).

PfACS knockdown induced a parasite growth phenotype, visible after the first 24h, where parasites became asynchronous and appeared unhealthy. It was noticed that most parasites had an amorphous appearance after 48h of knockdown and only a small number of parasites persisted after 72h. Growth assays also indicated a growth defect after PfACS knockdown, where 75% decrease of protein impaired parasite growth reaching only 0.5% parasitemia compared to 5% parasitemia in the controls. Additionally, the milder knockdown inhibition of 30% still impaired proper parasite growth. This underscores that PfACS is essential for parasite growth and development, in agreement with previously predicted experiments using piggyback insertion mutagenesis (41).

We also tested an ACS inhibitor that was reported to be effective against ACSS2. Considering the similarity between the catalytic sites of these two enzymes, we wondered if PfACS inhibition was also possible. When applying different concentrations of iACS, a growth phenotype very similar to the PfACS knockdown, and this effect was dose-dependent. The IC_50_ of iACS was 30.14 μM and the application of liposome encapsulation using multilamellar nanovesicles led to a 5.5-fold lower IC_50_, comparable to what was achieved in the study by Fotoran et al. (27). We also tested the effectiveness of iACS against *P. berghei* in the murine model and observed a significantly decreased parasitemia. This was expected due to the similarity between PbACS and PfACS. However, a relatively high dosage was needed to achieve even partial parasite clearance. Perhaps a modification of iACS structure may improve its interaction with PfACS, being then also more selective for the plasmodial ACS variant compared with human ACSS2.

PfACS knockdown led to significant decrease in epigenetic acetylation marks H3K14Ac and H3K9Ac, but not H4Ac. The same was observed after iACS treatment in a dosage-dependent manner, indicating that iACS is specific. This suggests that not all acetylation marks are influenced by PfACS. The presence of H3K4me3, an activation methylation mark that co-occupies many loci with H3K9Ac, was also tested and this modification was not affected by ACS inhibition. H3K9Ac is an activation mark that is spread across the parasite genome throughout the intraerythrocytic cycle, where it co-localizes with highly expressed gene loci (42) and is known for its presence at activated *var* gene loci (43). Mews and colleagues observed that ACSS2 binds to chromatin in CAD neuronal mouse cells. There, its genome location correlates positively with H3K9Ac presence while H3K9Ac without ACSS2 co-localization was observed in genes with less mRNA abundance (26). In *P. falciparum*, both knockdown and inhibition of PfACS led to a significant decrease of this mark, indicating that there may be a relation between them, but whether this protein in fact co-localizes with H3K9Ac marks remains to be studied. H3K14Ac is an activation mark and is also very abundant across the parasite genome where it reaches its maximum at ring stages 0-16 hours post infection (44). Interestingly, this correlates with PfACS presence in the nucleus, and among the tested marks this was the most affected acetylation mark by both PfACS knockdown and inhibition. It seems that nuclear PfACS directly contributes to H4K14 acetylation.

The H4Ac marks were also tested using an antibody that is reactive against different acetylated lysins in Histone 4. H4Ac has been described as occupying loci of active and poised genes and is predominantly found in sparsely expressed genes (42). Surprisingly, this mark was not significantly affected by PfACS knockdown or inhibition in *P. falciparum*, but in *P. berghei* a strong decrease was observed in qualitative immunoblots. Further quantitative analysis remains to be done to confirm this result. We observed a strong decrease in H3K9Ac, H3K14Ac and H4Ac acetylation marks in *P. berghei* treated with iACS, which indicates that similar to PfACS PbACS is an extremely important source of acetyl-CoA for protein acetylation. However, this protein has not yet been characterized in this species.

*P. falciparum* acetyl-CoA levels are maintained mostly by glucose metabolism, from glucose-derived pyruvate that undergoes BCKDH and hypothetically from acetate conversion by PfACS (24). Experiments measuring intracellular acetyl-CoA levels confirmed the importance of PfACS for providing acetyl-CoA. Unfortunately, the acetyl-CoA levels of parasites treated with iACS samples could not be determined by the Acetyl-CoA Assay Kit used in this work.

To establish a role of PfACS in *var* gene expression, a transcript profiling was performed using parasites selected to express CD36-binding PfEMP1-encoding *var* genes. The control transcribed successfully two CD36-binding PfEMP1 *vars*, PF3D7_1240600 and PF3D7_0412400, and PF3D7_0420900 at higher levels compared to the remaining *var* genes. Upon knockdown, a significant decrease of the steady state levels of the corresponding transcripts was observed. A very similar *var* transcript profile was observed in parasites treated with iACS. Possibly, inhibition of PfACS and decreased acetylation at active *var* loci occurred and this in turn led to decreased transcript levels and less PfEMP1. It is possible that the temporary decreased availability of Acetyl moieties interferes with the epigenetic memory of *var* transcription. This was tested by inducing PfACS knockdown or inhibition during one reinvasion cycle and subsequent outgrowth. As a result, the same *var* transcript profile as before PfACS knockdown or inhibition was observed. Only one single transcript returned significantly more abundant than the control upon iACS treatment, while the remaining *var* transcripts did not differ from the controls. In the case of PfACS knockdown, we observed a slight difference compared with the controls. The same set of *var* genes were found after knockdown, with the difference that they appeared in a significantly higher abundance than the controls. We also observed that the *var* transcript PF3D7_0632500, a gene known as one of the frequently expressed *var*s in *P. falciparum in vitro* cultures (45), became activated. This stronger transcription of *var* genes might be a compensatory effect resultant from PfACS knockdown that led to an up-regulation of this protein. Considering that the *var* transcript profile was not profoundly altered, PfACS seems not to play a direct role in the epigenetic memory of *var* gene transcription. We hypothesize that this protein acts in complex with histone-modifying proteins, providing the substrate for acetylation, but other proteins are responsible for regulating *var* locus activation and transcriptional memory.

It was also tested if the decrease of overall *var* transcripts under PfACS knockdown had consequences in the cytoadherence phenotype of IRBC on CHO cells expressing human CD36 receptors. In fact, a loss of the cytoadherence phenotype in PfACS knockdown parasites and in iACS treated parasites was observed, suggesting that *var* genes were de-activated during PfACS depletion, leading to less PfEMP1 expression and consequently less cytoadherence to CD36 receptors. In line with this hypothesis is the observation that PfACS and consequently *var* transcription returned to its previous levels, the cytoadherence phenotype was re-established. Parasites treated with iACS became more adherent to the CHO-CD36 cells compared with the controls, but there was no difference after PfACS knockdown. This phenomenon may be related to a compensating effect caused by PfACS inhibition.

PfACS was recently identified in complexes near *var* loci (30). In order to assess additional regions in the genome where PfACS interacts, ChIP-seq analyses were performed. PfACS clearly binds to chromatin and a peak calling analysis retrieved a list of 3002 different genes enriched in chromatin immunoprecipitation. Among these genes were many regulatory proteins, involved in central metabolic pathways, gene expression regulation and virulence multigene families, including *rifin, stevor, var, phist, surf* and *fikk*. Different from other organisms, *P. falciparum* is known to maintain a transcription permissive state of the chromatin most of the time (46). Thus, PfACS may be essential for euchromatin maintenance due to its importance in donating acetyl-CoA for histone acetylation, a mark that is mainly related to activation, and naturally this protein will interact with the chromatin surrounding genes important for parasite survival. We found that this protein mainly interacts in exonic regions across the whole parasite genome. Interestingly, most of the genes where PfACS was found interacting in intronic regions are classified in PlasmoDB as pseudogenes, and the reason for this is elusive. Among them are different pseudogenes from multigenic gene families such as *rifin, stevor, var, phist, surf* and *fikk*.

Among multigenic families enriched in the analysis, 38 PHIST genes and 11 PHIST pseudogenes were found. PHIST proteins are encoded by a large family of 89 genes. Although these proteins are known to be exported and have been linked to RBC remodeling and pathogenicity (47), the majority of them has not yet been characterized. PHIST proteins have different locations, including the IRBC surface (Yang et al., 2020), Maurer’s Clefts (49) and extracellular vesicles (50). Also, many members of this family have been found to be essential for parasite survival (41, 51). The PHISTb domain-containing RESA-like protein 1 (PF3D7_0201600), found in our analysis, showed to mediate specific interactions with PfEMP1 and to be somehow involved in *var* gene expression regulation (52). Another PHIST that might be regulated by PfACS is the Lysine-rich membrane-associated PHISTb protein (PF3D7_0532400), one of the ATS binding PHISTs that is localized at IRBCs knobs, that plays a role in cytoadherence to CD36 receptors (53).

Mews and colleagues identified that ACSS2 interacts with chromatin in CAD neuronal mouse cells. These authors also found many enriched peaks in common with H3K9Ac marks peaks, indicating that the presence of this protein colocalize with this activation mark (26). Possibly, the same result might be found in *P. falciparum*, however, a parallel ChIP-seq of PfACS, H3K9Ac and other acetylation marks must be performed to address this question.

Our ChIP-seq analysis using ring stage parasites revealed PfACS enrichment in 7 *var* gene loci and 13 pseudogenes. For this experiment, a lineage with no previous cytoadherence-selection was used. Therefore, *var* gene expression is expected to be mixed as it normally is in *in vitro* cultures. Interestingly, among the few genes where PfACS was identified at their promoter regions is the PfEMP1 PF3D7_0800100, which was identified together with PF3D7_0421300 to be the most commonly activated genes after *var* switching (54–56). Also, this locus is usually one of the most transcribed in long-term *in vitro* cultures (45, 57). More recently, Bryant and colleagues described a protein complex at *var* gene activated promoters that includes, besides known factors such as H2A.Z, SIP2 and HP1, the putative chromatin remodeler PfISWI, PfACS and an ApiAP2 transcription factor (30). These results corroborate the observation that PfACS interacts with *var* loci.

Given its influence in multiple aspects of the intraerythrocytic life cycle of the parasite PfACS or proteins that may intermittently interact with it may be excellent targets for drug development. In addition, structure-guided modifications of already existing drugs may prove strong candidates for malaria therapy.

## Material and Methods

### Transfection plasmid construction

For single site homology recombination, a 1115 basepair fragment from the 3’ end of the *PfACS* gene excluding the stop codon was generated by standard PCR using the oligonucleotides *ACS* Forward 5’-agatctCAACATATCCAGATTGTGGTAG-3’ and *ACS* Reverse 5’-ctgcagcTTTCTTAATTTCAATATGCTTTAAC-3’, Elongase Enzyme Mix (Invitrogen) and genomic DNA from *P. falciparum* strain NF54. The fragment was amplified, agarose gel-purified, ligated into the pGEM T Easy vector (Promega) and transformed in DH10B chemo-competent cells. Resulting clones where Sanger-sequenced and a 100% identical fragment was subcloned from the corresponding pGEM clone via restriction enzymes PstI and BglII. The fragment was ligated into the p_GFP-HA-DD24-2A-BSDglmS vector, generating pACS-GFP-HA-DD24-2A-BSDglmS. Once added, the fragment became fused to the GFPHA tag followed by the degron DD24 (35). This domain is followed by the 2A skipping peptide and Blasticidin S deaminase, which confers resistance to Blasticidin after homologous recombination in the *ACS* locus, and additionally adds the glmS ribozyme. The homology region allows the single crossover recombination and consequently the integration of the fragment and its following features to the parasite genome. The vector also contains the hDHFR cassette, which provides resistance to WR99210 drug (a gift from Jacobus Inc., USA) for the initial positive selection of transfected parasites.

### P. falciparum culture

*P. falciparum* NF54 parasites were cultivated at 4% hematocrit in human B+ blood in RPMI-HEPES supplemented with 0.25% Albumax 1 (Invitrogen) and human donor plasma type B+. Ethical clearance for the use of human blood and plasma was obtained from the local ethics commission at the Institute for Biomedical Sciences (CEPSH-ICB/USP, protocol No. 874/2017). The medium was changed every day or every two days and parasites were fed with fresh blood every 4 days or more often depending on the culture parasitemia. When tightly synchronized cultures (4-6 h window) were needed, mature stage parasites were density-floated using Voluven 6% (Fresenius Kabi, Campinas, Brazil) using a protocol described by Lelièvre and colleagues (58). For ring stage parasite purification, cultures were treated for 10 min at RT with 20 volumes of a 5% Sorbitol solution (59).

### Parasite Transfection

The protocol for plasmid transfection was described in detail by Hasenkamp and colleagues (60). Briefly, the protocol consists of the electroporation of fresh RBCs with 40 µg of the highly purified plasmid. In parallel, an NF54 parasite culture is Voluven-treated (see above) to yield enriched schizont stages at 40-80% parasitemia. These were then transferred to the culture bottle containing exclusively electroporated RBCs. Culture media of transfected parasites were changed daily and the WR99210 drug was added on the second day after transfection at 2.5 nM. Approximately ∼20 days after transfection, WR99210 resistant parasites appeared and backups were frozen. Then, Blasticidin S at a final concentration of 2.5 µg/mL, was added to the culture leading to the positive selection of parasites with locus-integrated plasmids, eliminating episome-carrying parasites. After ∼17 days, parasites became again detectable in blood smears. Conventional PCRs using primers targeting a region upstream (*ACS* check Forward 5’-AGGTTGGGTTACCGGACATAC-3’) of the homology region used for cloning and a reverse primer targeting the HA encoding sequence (3xHA Reverse: 5’-AGCGGCATAATCTGGAACATCGTAC-3’) were performed to confirm the success of genome integration. In addition, PCRs using a primer targeting unmodified locus (*ACS 3’UTR* Reverse 5’-TGTGGGGCTTCAAAATATGAG-3) were performed to check the complete absence of unmodified *PfACS* loci.

### Panning Assay

After NF54::ACSGFPHADD242AbsdglmS mutant line generation, two panning assays were performed using the protocol established by Gölnitz and colleagues (28). The objective was the selection of parasites transcribing predominantly one or a few *var* genes expressing CHO-CD36-binding PfEMP1 phenotype. For this, parasites were selected for cytoadherence by letting trophozoite/schizont stage parasitized iRBC adhere to confluent CHO-CD36 cells. Briefly, Voluven-floated trophozoite stage iRBC (parasitemia of 5⨯10^7^-10^8^) were layered over confluent CHO-CD36 cells in RPMI/10% human plasma at pH 6.8. The iRBC were incubated over the cells for one hour at 37°C, being softly stirred every 15 minutes. After 1h, the non-adherent iRBCs were removed and the remaining adherent iRBCs were gently washed three times with RPMI at pH 6.8. The cytoadherent parasites were detached using complete culture medium at pH 7.2-7.4 and returned to normal culture conditions. This process was repeated twice.

### Quantitative Cytoadherence Assay

After NF54::ACSGFPHADD242AbsdglmS were selected to express specific *var* genes and were therefore cytoadherent to CHO-CD36 cells, we established the following conditions to measure cytoadherence: PfACS ON (Control), PfACS OFF (ACS Knockdown), iACS (Presence of ACS inhibitor) and PfACS Recovery. Cytoadherence assays were then carried out using the protocol described above (28). After the three washing steps with RPMI pH 6.8, fifteen pictures for each condition were taken using EVOS FL Digital Inverted Microscope (AMG). Experiments were carried out in 6-well plates in three biological replicates. The number of adherent iRBCs and CHO cells were calculated using ImageJ software (NIH).

### Knockdown Assays

The mutant line NF54::ACSGFPHADD242AbsdglmS permits the knockdown of PfACS at two levels. At the transcriptional level, the glmS ribozyme element induces it self-cleavage in the presence of Glucosamine and, as it is located at the 3’ end of the mRNA of interest, the transcript polyA tail is removed, which turns the transcript unstable and susceptible to degradation (61). At the protein level, the destabilizing domain DD24 maintains protein stability only in the presence of Shield-1, and, when this compound is removed, the protein becomes unstable (62). The knockdown assays were conducted by using NF54::ACSGFPHADD242AbsdglmS parasites continuously maintained in the presence of 0.5 µM Shield-1 as control group, Shield-1 withdrawal as second knockdown group, and Shield-1 withdrawal plus addition of 2.5 mM Glucosamine/HCl to the culture medium as third knockdown group.

### *In vitro* ACS Inhibition Assays

1-(2,3-di(Thiophen-2-yl)quinoxalin-6-yl)-3-(2-methoxyethyl)urea (CAS 508186-14-9 – MCE-MedChemExpress) is an Acetyl-CoA Synthetase 2-specific and potent inhibitor, tested in human (25) and mouse cells (26) . NF54 WT parasites were submitted to different doses of ACS inhibitor (iACS), ranging from 0 (DMSO used as control) to 100 µM. Parasites were cultivated at 0.15% parasitemia and 2.5% hematocrit in 96-well plates, each well containing 200 µL of parasite culture, and submitted to eight 2-fold iACS dilutions and its IC_50_ was determined. In an attempt to improve inhibitor delivery, we performed iACS compound encapsulation into multilamellar liposomes as described by Fotoran and colleagues (27) and the IC_50_ assay was carried out as described above. IC_50_ analyses were performed in GraphPad Prism 5 software.

### *In vivo* ACS Inhibition Assays

To test iACS efficacy against *P. berghei* in murine model, five CL57BL/6 mice per treatment group were infected intraperitoneally with 10^5^ *P. berghei* ANKA strain GFP-expressing parasites. Starting at post-infection day 2, mice were treated intraperitoneally for three consecutive days, where the control mice group was injected with DMSO and the iACS mice group received 35 μM of iACS in DMSO. Parasitemia was calculated from blood sample of each mouse by flow cytometry (63) of 30000 RBCs and checked by stained blood smears. Also, using two extra mice, one control DMSO-treated and one iACS 30 μM for one day, total protein extracts from mice blood parasites were submitted to western blotting. Ethical clearance for this experiment was granted by the local Ethics Commission for the Use of Animals in Research (CEUA protocol 015, published 19.3.2013).

### Western Blotting

Parasite cultures were collected, 0.1% Saponin lysed, washed 3x in PBS and diluted in SDS cracking buffer to obtain total protein extracts. Samples were then separated in 10% SDS polyacrylamide gels and transferred to Hybond C nitrocellulose membranes (GE Healthcare). After blocking with 5% non-fat milk in PBS/0.05% Tween, membranes were incubated with primary antibodies overnight incubation in 1% non-fat milk PBS/0.05% Tween20. Blots were washed 3x during 15min in PBS/0.05% Tween and incubated for 1h with the secondary antibodies in 1% non-fat milk PBS/0.05% Tween20. After triple PBS/0.05% Tween washings, blots were incubated with WesternPico Super signal substrate (Pierce/Thermo) for detection and chemoluminescence was visualized with X-Ray films (Hyperfilm, GE Healthcare). The following antibodies were used in this work: Anti-HA (Cell Signaling), Anti-ATC (in-house produced in mice), Anti-H3 (Millipore), Anti-H3K9Ac (Millipore), Anti-H3K14Ac (Millipore), Anti-H4Ac (Millipore), Anti-H3K4me3 (Millipore), Anti-TPK (donated by Dr. C. Wrenger) and Anti-β-Tubulin (Cell Signaling). Anti-mouse (KPL) and Anti-rabbit (KPL) coupled to peroxidase were used as secondary antibodies. The signal intensities were measured using the ImageJ program (NIH) and quantitative analysis was performed in GraphPad Prism 5 software.

### Fluorescence Microscopy

Mutant parasites were submitted to fluorescence microscopy to detect GFP expression. For that, parasites were fixed as described in by Tonkin and colleagues (64). After fixation, parasites were incubated in PBS/Saponin 0.01% and DAPI at a final concentration of 2 µg/mL at 37°C for 1h. After that, parasites were washed 3x with PBS/Saponin 0.01% and incubated with the membrane marker Wheat Germ agglutinin (WGA) Texas RedTM-X conjugate (Invitrogen) in PBS/Saponin 0.01% for 20 minutes at 37°C. Parasites were then washed 3x with PBS/Saponin. Images were obtained with a fluorescence microscope (DMI6000B/AF6000, Leica, Germany) connected to a digital camera system (DFC 365 FX, Leica, Germany) and processed by Photoshop 5 software.

### RNA Extraction and cDNA Synthesis

Parasites were synchronized in ring stage (2-8 h p.i.) and ∼700 µL at ∼6% parasitemia of the infected blood was lysed in PBS/Saponin 0.1%, washed 1x with PBS, resuspended in 100 µL of PBS and further in 1000 µL TRIzol (Life Technologies). RNA was then extracted following the manufacturer’s instructions. The final RNA pellet was resuspended in 20 µL nuclease-free water. For reverse transcription, 5 μg of total RNA was digested for 30 min at 37°C by DNAse I (Fermentas) enzyme. RNAs were then used for cDNA synthesis using SuperScript IV Reverse Transcriptase (ThermoFisher Scientific) and random hexamer oligonucleotides following the manufacturer’s instructions. RNA quality was checked by electrophoresis in Agarose 0.8% TBE gels.

### *Var* Gene Profiling by RT-qPCR

RT-qPCR assays were performed using 5x HOT FIRE Pol EvaGreen qPCR Mix Plus (Solis Biodyne) in a QuantStudio 3 Real-Time PCR System (ThermoFisher Scientific) machine. Relative transcript quantities of all 3D7 *var* genes (see a list of oligos used for RT-PCR in Supplementary Table 3) were calculated by the 2^-ΔCt^ method (65) using plasmodial Seryl tRNA Ligase (“K1”) transcript as the endogenous control.

### ChIP-seq Library Generation and Analysis

NF54::ACSGFPHADD242AbsdglmS ring stage synchronous cultures were harvested, crosslinked with 1% formaldehyde and incubated for 15 minutes at 37 °C with agitation. Crosslinking was stopped by Glycine addition at a final concentration of 0.125 M and for 5 minutes of incubation on ice. Samples were washed 3x with PBS followed by 10 minutes incubation with saponin 0.10 %. Following this, parasites were centrifuged at 4000 rpm, 4 °C for 15 minutes and pellets were washed 3x with PBS. Samples were resuspended in LowRIPA buffer, transferred to a dounce homogenisator and NP-40 at a final concentration of 0.25% was added. After 100 pulses, samples were centrifuged for 10 min at 4°C 14000 rpm. Then, chromatin was immunoprecipitated using the MultiMACS HA Isolation Kit (12×8, Miltenyi Biotech) following manufacturer’s instructions. Briefly, pellets were resuspended in kit lysis buffer and anti-HA monoclonal antibody conjugated to µMACS MicroBeads were added to the samples. After 30 min resting on ice, samples were placed in the µMACS™ Separator for another 30 min. Samples were then washed 4x and immunoprecipitated protein/DNA was eluted. Importantly, whole cell extracts not submitted to immunoprecipitation, were withdrawn from the samples before immunoprecipitation and treated similarly during all steps. The DNA present in the samples was extracted with DNA extraction buffer (0.1 M EDTA pH 8, 1% SDS and 200 µg/mL proteinase K) overnight at 65°C. DNA was then extracted by Phenol-Chloroform extraction. We next performed the library preparation for sequencing using TruSeq ChIP Library Preparation Kit (Illumina), following the manufacturer’s instructions. Quality of the libraries was tested using a Bioanalyzer 2100 (Agilent). Sequencing was carried on an Illumina NextSeq® 500/550 system as a 75 bp single-end run. Two biological replicates plus input of each sample were sequenced and analyzed. Low quality region (<Q20) elimination of reads was done by the program Sickle (Joshi, https://github.com/najoshi/sickle) using the parameters -q 20 for minimum quality and -l 100 for minimum length. Read alignments were performed using the software Burrows-Wheeler Aligner (BWA) at default parameters (66). For the alignment and subsequent steps, *P. falciparum* 3D7 (GenBank assembly accession: GCA_000002765.2) was used. The annotation file Plasmodium_falciparum.ASM276v2.45.gff3 is available in the following link: ftp://ftp.ensemblgenomes.org/pub/protists/release-49/gff3/plasmodium_falciparum. Peak calling analysis of read alignments was performed by the software MACS2 (Zhang et al., 2008) using macs2 callpeak and ‘-g 2e7 –nomodel --extsize 75 --keep-dup auto’ parameters. Fraction of reads in peaks (FRiP), Normalized Strand Cross-correlation coefficient (NSC), and Relative strand cross-correlation coefficient (RSC) were obtained by NGS tools available in the following link: https://github.com/mel-astar/mel-ngs/tree/master/mel-chipseq/chipseq-metrics. ENCODE recommended values for NSC, RSC and FRiP are >1,05; >0,8 and >0,01 respectively (68). Peaks found in regions until 1.6 kb and 60 kb from chromosomes extremities were considered telomeric and sub-telomeric, respectively. Peaks found -/+ 1kb from transcription start sites were considered promoter region-localized.

### Acetyl-CoA Assay

In order to determine the cellular amount of Acetyl-CoA in parasites in the under knockdown or not or during inhibition of PfACS, the following groups of treated parasites were analyzed: NF54::ACS-GFP-HA-DD24-2A-bsd-glmS parasites cultivated in the presence of Shield-1 (Control group), and for 24 h in the absence of Shield-1 plus Glucosamine (Knockdown group); NF54 WT parasites treated with vehicle (DMSO, Control group), or 30, 40 and 50 µM iACS for 24h. IRBC were lysed by Saponin 0.1% and washed 3x in PBS. Pellets were resuspended in Cell Lysis Buffer (20 mM Tris (pH 7.5), 150 mM NaCl, 1 mM EDTA, 1 mM EGTA, 1% Triton X-100, 2.5 mM sodium pyrophosphate and 1 mM β-Glycerolphosphate) and sonicated 4x for 5s. Samples were then deproteinized by PCA precipitation. Briefly, perchloric acid was added to each sample at a final concentration of 1M, vortexed, centrifuged at 10,000g for 5 min and neutralized with 2M KOH until each sample pH reached 6-8. After that, samples were centrifuged at 10,000g for 5 min and the Acetyl-CoA measurement was performed in a 96-well plate using the Acetyl-Coenzyme A Assay Kit (Sigma-Aldrich) following the manufacturer’s instructions. Fluorescence intensity was measured in the microplate reader CLARIOstar Plus (BMG Labtech) using λ_ex_=535/ λ_em_=587.

### Cytoplasm and Nucleus Fractioning

The cell compartments cytoplasm and nucleus were fractionated in order to localize PfACS. For that, parasites were synchronized in ring, trophozoite and schizont stages, lysed with PBS/Saponin 0.1% and washed 3x with PBS. Cell-fractionation of parasites was done using the protocol as described by Oehring and colleagues (69). Pellets were gently resuspended in the hypotonic Cytoplasm Lysis Buffer (20 mM HEPES (pH7.9), 10 mM KCl, 1 mM EDTA, 1 mM EGTA, 0.65% NP-40, 1 mM DTT and protease inhibitors), put on ice for 5 min and centrifuged at 2,000g for 5 min at 4°C. These supernatants were saved as the cytoplasmic fractions. Pellets were gently washed 7x with Cytoplasmic Lysis Buffer and then resuspended in Low Salt Buffer (20 mM HEPES (pH7.9), 0.1 M KCl, 1 mM EDTA, 1 mM EGTA, 1 mM DTT and protease inhibitors). These samples were saved as parasite nuclear fractions. After that, all samples were submitted to western blotting.

## Supplementary Material: Figures

**Supplementary Figure 1.**
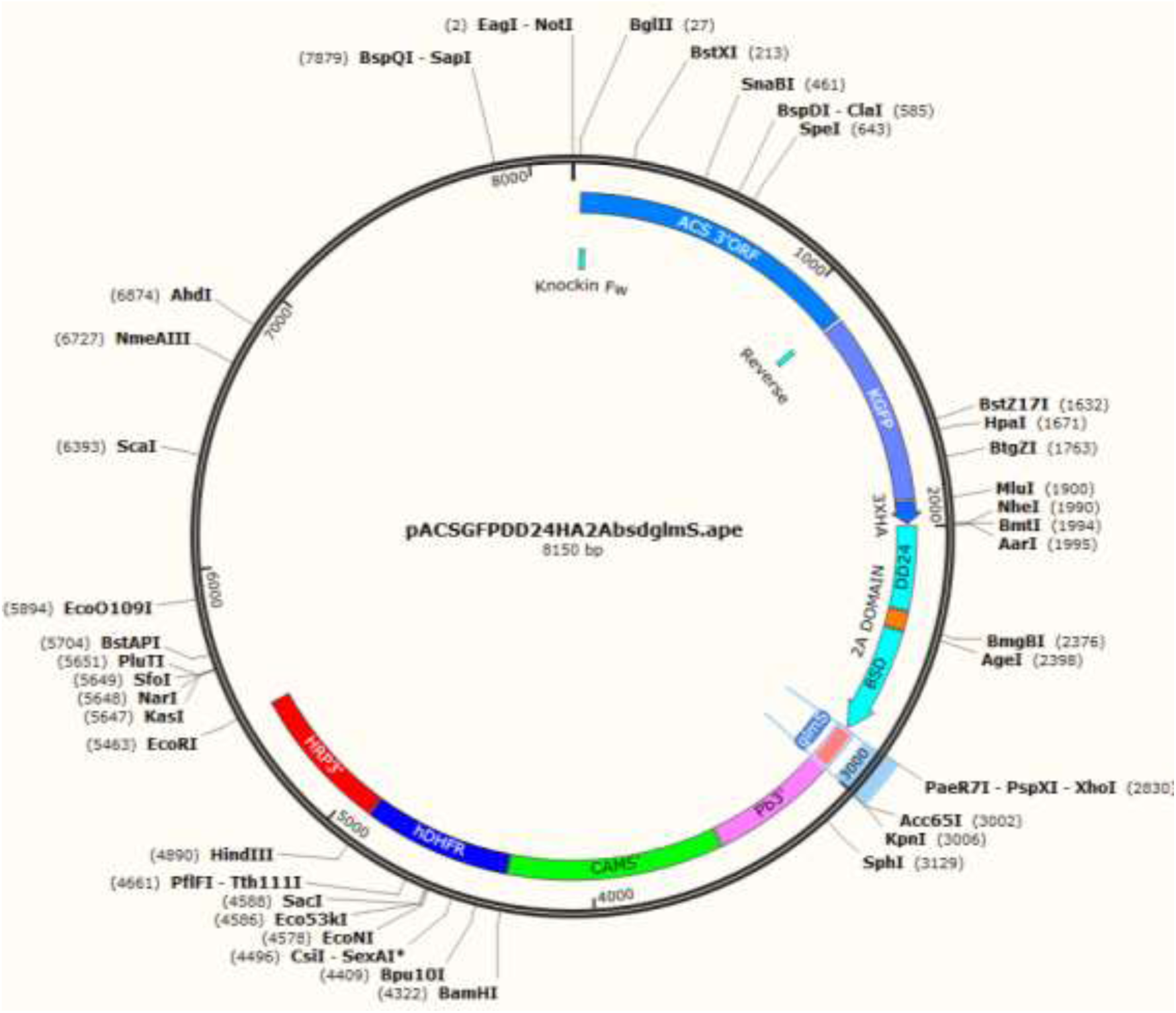
Map of the pACS-GFP-HA-DD24-2A-BSD-glmS vector construct. The vector sequence is available upon request.

**Supplementary Figure 2.**
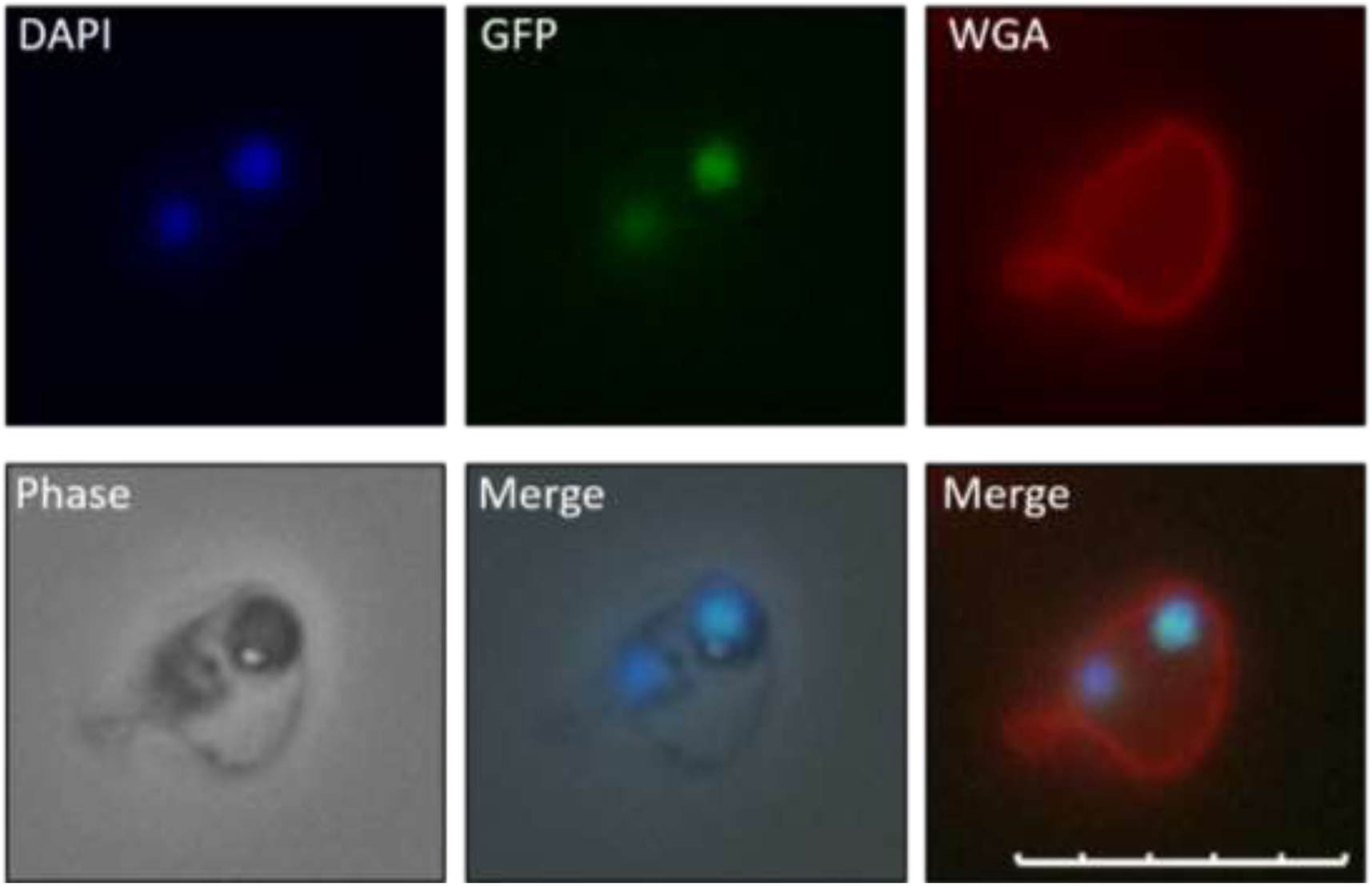
Fluorescence microscopy of ring stage *P. falciparum* parasites exhibiting GFP expression. DAPI was used as parasite nuclear marker and WGA as erythrocyte surface marker. Scale bar: 10 µm.

**Supplementary Figure 3.**
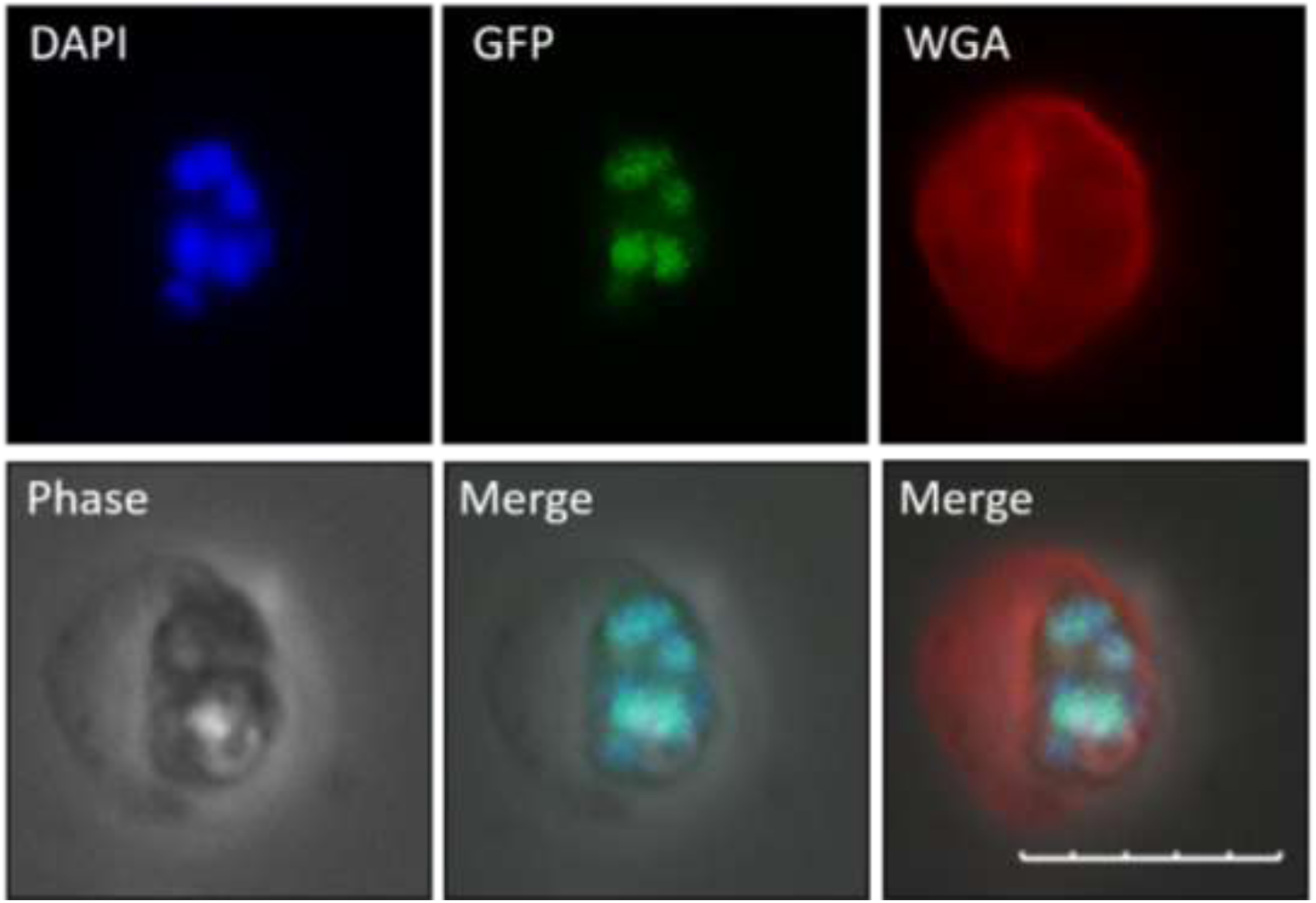
Fluorescence microscopy of late trophozoite stage *P. falciparum* parasites exhibiting GFP expression. DAPI was used as parasite nuclear marker and WGA as erythrocyte surface marker. Scale bar: 10 µm.

**Supplementary Figure 4.**
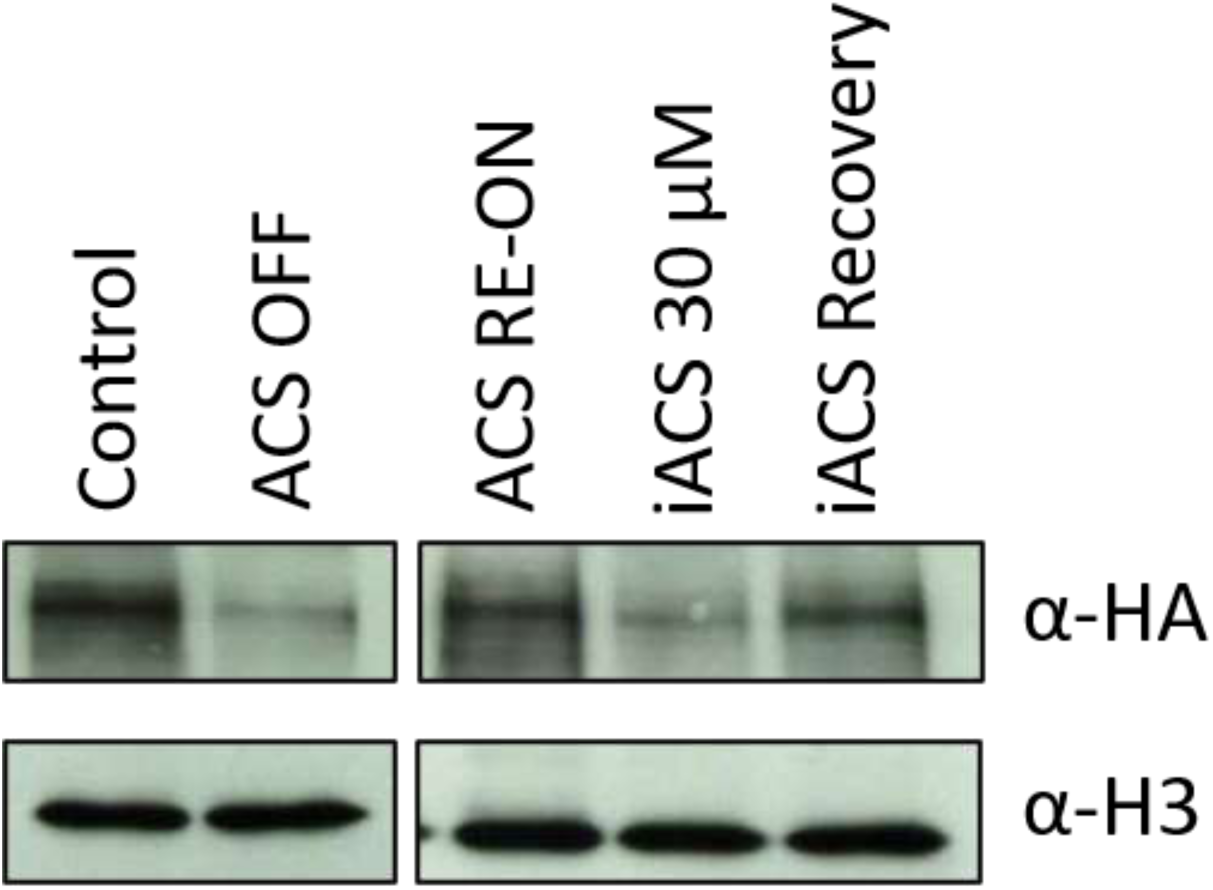
Western blotting indicating PfACS knockdown (ACS OFF) and, after knockdown treatment retrieval, protein re-establishes protein synthesis (ACS RE-ON). Also, presence of iACS inhibitor and treatment retrieval can be seen.

